# Development of a mouse salivary gland-derived mesenchymal cell line for immunological studies of murine cytomegalovirus

**DOI:** 10.1101/2022.03.03.482882

**Authors:** Timothy M. White, Brent A. Stanfield, Cassandra M. Bonavita, Jared S. Rudd, Rhonda D. Cardin

## Abstract

The salivary glands are a crucial site of replication for human cytomegalovirus (HCMV) and its murine counterpart, murine cytomegalovirus (MCMV). Studies of MCMV often involve the use of BALB/c strain mice, but most *in vitro* assays are carried out in the NIH 3T3 cell line, which is derived from Swiss Albino mice. This report describes a BALB/c-derived mouse salivary gland cell line immortalized using the SV40 large T antigen. Cells stained positive for PDGFR1 and negative for E-cadherin and PECAM-1, indicating mesenchymal origin. This cell line, which has been named murine salivary gland mesenchymal (mSGM), shows promise as a tool for *ex vivo* immunological assays due to its MHC haplotype match with the BALB/c mouse strain. In addition, plaque assays using mSGM rather than NIH 3T3 cells are significantly more sensitive for detecting low concentrations of MCMV particles. Finally, it is demonstrated that mSGM cells express all 3 BALB/c MHC class I isotypes and are susceptible to T cell-mediated *ex vivo* cytotoxicity assays, leading to many possible uses in immunological studies of MCMV.

## Introduction

Several medically significant viruses, including Epstein-Barr virus, (EBV) human cytomegalovirus (HCMV), and, as recently reported, severe acute respiratory syndrome coronavirus 2 (SARS-CoV2) utilize the salivary glands as a site of replication and spread [1,2]. *In vitro* studies of salivary gland cells frequently focus on acinar epithelial cells, which are responsible for the production and secretion of saliva [3]. Additionally, the acinar epithelium is an important site of replication for HCMV and murine cytomegalovirus (MCMV). Virus secreted in saliva derived from acinar tissue is considered critical to the horizontal transmission of both viruses [3]. In addition to acinar epithelial cells, murine salivary glands consist of endothelial cells, which line blood and lymphatic vessels, as well as mesenchymal cells, which provide supporting roles in development and healing. Mesenchymal cells, which include fibroblasts, myofibroblasts, and other stromal cells are found in tissues throughout the body, and may serve as a reservoir of latent MCMV and HCMV [4,5].

Mouse embryonic fibroblasts (MEFs), most often the NIH 3T3 cell line, are commonly used for *in vitro* MCMV studies and are extensively used for production of virus stocks and plaque assays for determining virus titers. NIH 3T3 cells [6] are the prototypical mesenchymal cell line for most *in vitro* mouse studies, and additionally are a useful tool for many assays across varied disease models. However, NIH 3T3 cells are derived from the Swiss-Webster albino strain of mice [6]. Therefore, cell surface molecules, including MHC isotypes, are not compatible with many *ex vivo* studies in the standard BALB/c mouse model of MCMV infection.

One assay used to investigate latent herpesviruses, including herpes simplex and MCMV, is *ex vivo* culture of latently infected tissues, from which latent virus reactivates and begins lytic replication [7–9]. After explantation of tissues (e.g., salivary gland, spleen, liver, and lung), fibroblast-like cells migrate out of tissue fragments and form a monolayer. Our efforts to produce a salivary gland-derived cell line were motivated by this observation: during explant reactivation studies of MCMV, cytopathic effect (CPE) in migratory fibroblast-like cells was observed, suggesting viral infection and replication; however, the lineage of these cells and their permissiveness to MCMV infection and replication was unknown.

Here, we report on a salivary gland-derived mesenchymal cell line cultured from BALB/c ByJ mice and immortalized using SV40 large T antigen. This cell line, which has been termed murine salivary gland mesenchymal (mSGM), is permissive to MCMV. Analysis of surface markers confirmed mesenchymal lineage. Importantly, the mSGM cell line expresses all 3 isotypes of MHC found in BALB/c mice, allowing for cytotoxicity assays using CD8+ T cells with diverse epitope specificities, thus providing a useful tool for immunological studies of many pathogens, including MCMV.

## Results

### mSGM cells exhibit fibroblast-like morphology and markers of mesenchymal lineage

Salivary gland cells were isolated and cultured as described in the *Materials and Methods* section. Primary salivary gland cells were polymorphic, however, after 2-3 passages, cultures became uniformly dominated by contact-inhibited fibroblast-like cells. These cells could be maintained in culture for 8-10 passages before they began to lose their contact inhibition and form three-dimensional clumps. After 1-2 further passages, the primary cells no longer adhered to tissue culture-treated plates and culture was discontinued. In order to preserve the monolayer-forming fibroblast-like cells, primary salivary gland cells (passage 3) were transfected with a plasmid expressing the SV40 large T antigen and a stable subculture (designated mSGM) was produced as described. mSGM cells were grown to approximately 80% confluence and imaged, showing typical fibroblast-like morphology (Fig 1a). Next, mSGM cells were fixed and stained with antibodies against platelet endothelial cell adhesion molecule (PECAM-1), expressed on endothelial cells and leukocytes (Fig 1b); epithelial cadherin (E-cadherin), expressed on epithelial cells (Fig 1c); platelet-derived growth factor receptor beta (PDGFR1), expressed on mesenchymal cells (Fig 1d); and isotype control (Fig 1e). Cells were found to be strongly positive for PDGFR1 but negative for PECAM-1 and E-cadherin, indicating mesenchymal lineage [17].

**Figure 1.**
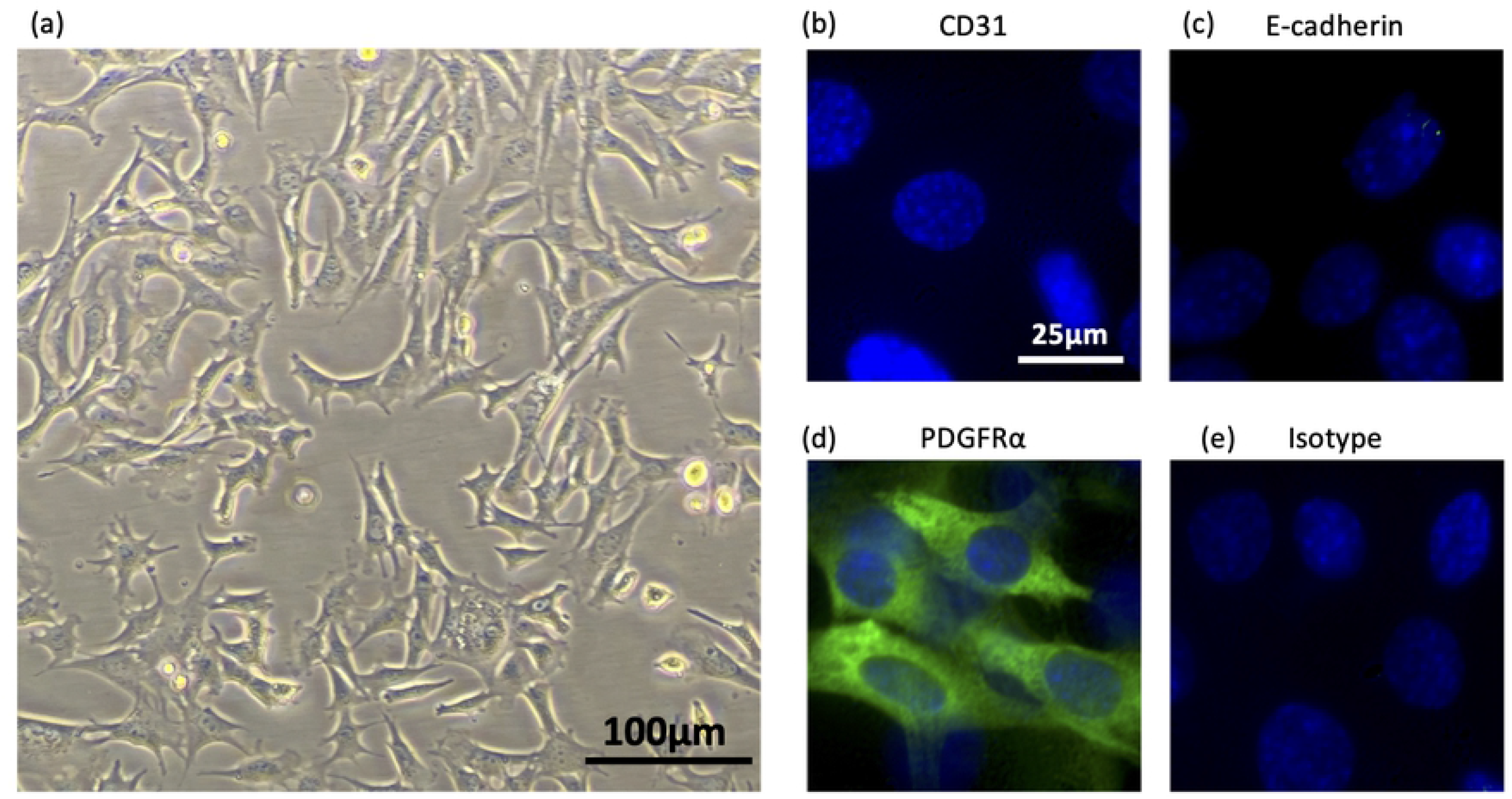
mSGM cells exhibit fibroblast-like morphology and markers of mesenchymal lineage. (a) mSGM cells were grown to subconfluence and live cells were imaged at 100x magnification. (b-e) mSGM Cells were stained for CD31, an endothelial cell marker (b); E-cadherin, an epithelial marker (c); PDGFR1, a mesenchymal marker (d); IgG2 isotype control (e).

### mSGM cells are permissive to MCMV infection and can be used for plaque assay

In order to determine whether mSGM cells remained permissive to MCMV infection after immortalization, single- and multi-step growth curves were conducted using wild-type MCMV (KP strain). MCMV replication in mSGM cells was found to be nearly identical to growth in NIH 3T3 fibroblasts at both low and high multiplicity of infection (MOI) (Figs 2a-b). No significant differences were found at any time point. We therefore concluded that mSGM cells are permissive to MCMV infection and yield quantities of virus similar to NIH 3T3 cells.

**Figure 2.**
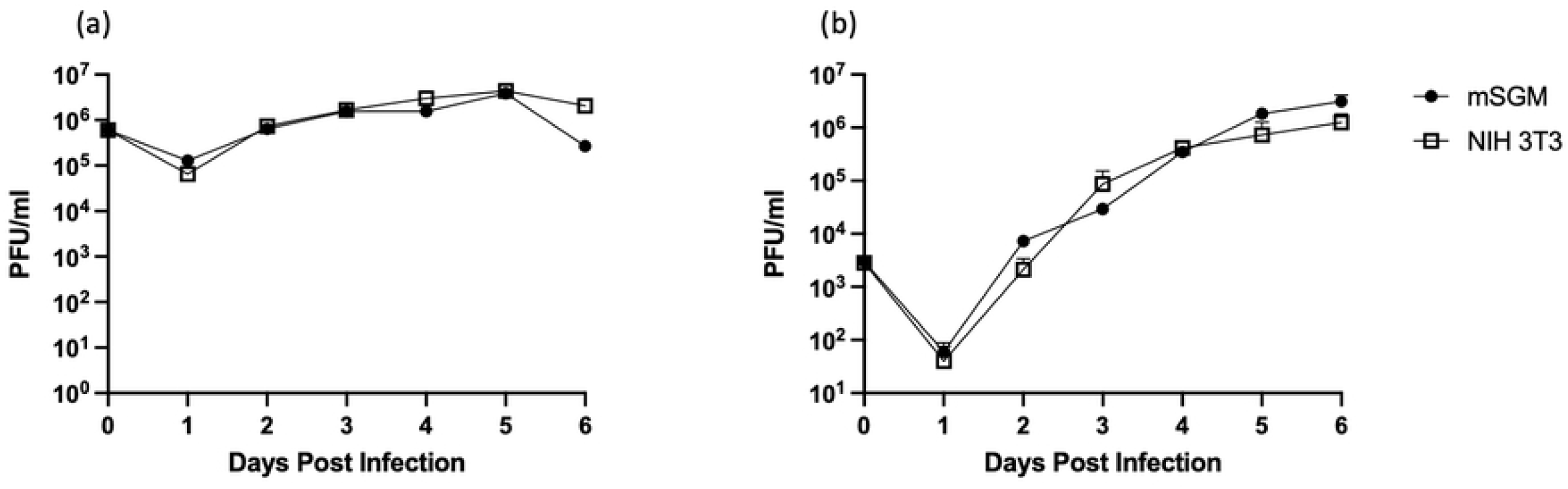
mSGM cells are permissive to MCMV infection and replication. mSGM cells and NIH 3T3 cells were infected with KP strain MCMV. (a) cells were infected at MOI of 2 for single-step growth. (b) cells were infected at MOI of 0.01 for multi-step growth. No significant differences were found at any time point (n=6).

Some cell lines, including the NIH 3T3 cell line, become less permissive to MCMV replication at higher passage numbers, necessitating frequent re-starting of cultures. In order to determine the stability of the mSGM cell line at higher passages, cells were passaged 2-3 times weekly for 6 months. After 40 passages, morphology had not changed, and plaque efficiency remained comparable to low-passage mSGM cells. After 60 passages, however, the mSGM cells were found to divide too rapidly for efficient plaque formation. Nevertheless, this represents a substantial improvement over NIH 3T3 cells, which in our hands are useable only to passage 18-20 before their permissiveness to MCMV declines. Additionally, mSGM cells were tolerant of freeze-thaw cycles in complete medium with 5% DMSO.

To determine whether mSGM cells could be used for plaque assays, studies using identical virus dilutions on NIH 3T3 cells or mSGM cells were performed, overlaid with identical medium, and stained at 5 dpi. Both cell types grew well under overlay medium and plaques could be visualized in both cell lines, although mSGM cells tended to produce larger plaques with more clearly defined borders (Fig 3a-b). Efficiency of plaque formation was compared between the two cell lines and mSGM cells were found to produce significantly greater numbers of plaques than NIH 3T3 cells when the same MOI of virus was used (Fig 3c). The differences in plaque number were more dramatic when the cultures were subjected to centrifugal enhancement [11]. Centrifugal enhancement, in which the 1-hour incubation at 37° is replaced with a 1-hour centrifuge step prior to overlay, resulted in >100-fold increase in plaque number over standard NIH 3T3 assay (Fig 3c).

**Figure 3.**
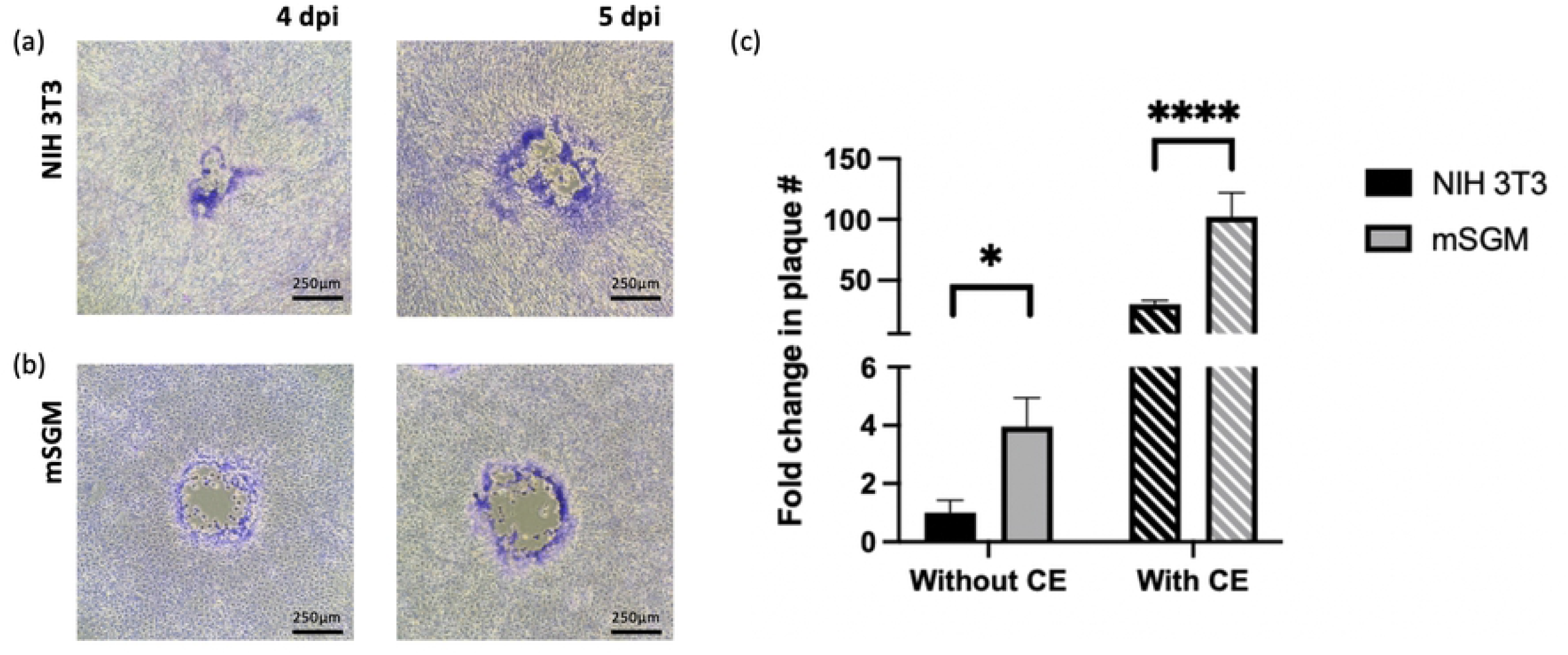
mSGM cells are suitable for highly sensitive MCMV plaque assay. (a-b) NIH 3T3 cells (a) and mSGM cells (b) were infected with 30 plaque forming units (pfu) (initially determined on NIH 3T3 cells) per well of a 12-well plate. After 4 (left column) or 5 days (right column) under overlay, plaques were stained with Giemsa. Images show representative plaques. (c) mSGM cell plaque numbers compared to NIH 3T3 cells when inoculated with identical pfu per well without centrifugal enhancement (without CE; solid bars) or with centrifugal enhancement (with CE; lined bars). Images are at 40x magnification on an inverted microscope. Significance is indicated as *, p<0.05; ****, p<0.0001 (n=6).

### mSGM cells express three isotypes of MHC class I

One of the key drawbacks to using NIH 3T3 cells for studies of MCMV is the difference in MHC haplotypes between Swiss Webster albino mice and other mouse strains, making NIH 3T3 cells unsuitable as antigen presenting cells for immunological studies using BALB/c mice. We thus investigated whether the mSGM cell line expresses MHC on its cell surface. BALB/c mice express 3 MHC class I isotypes: H-2D^d^, H-2K^d^, and H-2L^d^ [18]. In order to determine the expression levels of each isotype, mSGM cells were stained with antibodies against each of these MHC isotypes and found to express all 3 isotypes at high levels on the cell surface (Fig 4a-c). This suggested that mSGM cells would present a variety of peptide epitopes as seen in BALB/c mice *in vivo*.

**Figure 4.**
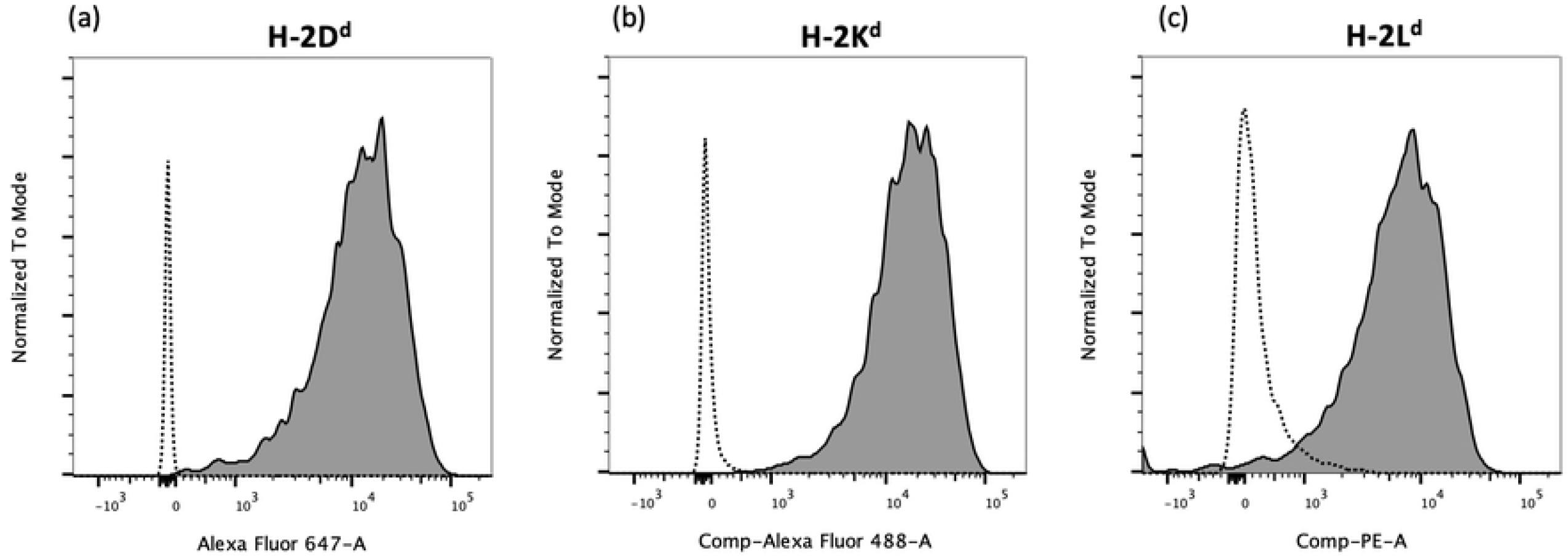
mSGM cells express 3 isotypes of MHC class I. mSGM cells were stained with antibodies against MHC class I H-2D^d^ (a), H-2K^d^ (b), or H-2L^d^ (c). Isotype controls are shown for comparison (dotted lines).

### mSGM cells are vulnerable to *ex vivo* MCMV antigen-specific T cell-mediated cytotoxicity

Besides MHC class I molecule expression on the cell surface, processing and presentation of viral protein-derived peptide epitopes is essential for T cell-mediated cytotoxicity. Therefore, mSGM cells were tested as antigen presenting cells for MCMV-specific T cell cytotoxicity. T cells were enriched from latently MCMV-infected or mock-infected BALB/c mice. An adenovirus vector expressing the MCMV immediate-early protein 1 (IE1), a major CD8+ T cell epitope (Ad5-MCMV_IE1) [19], or a matched adenovirus control (Ad5-control) which expressed only an irrelevant sequence, were used to introduce MCMV antigens. Uninfected mSGM cells were used as a secondary control. Enriched T cells were added at an effector:target (E:T) ratio of 6:1 and cytotoxicity was monitored using an Incucyte live-cell imaging system, similar to previous T cell cytotoxicity studies [20]. Apoptosis was quantified using NucView 488, a non-toxic substrate which fluoresces brightly after cleavage by caspase-3 [21]. T cell-mediated cytotoxicity was most apparent at 48 hours, which allowed time for rare epitope-specific T cells to encounter and interface with target cells.

As expected, uninfected mSGM cells and Ad5-control infected mSGM cells, neither of which produces MCMV-associated epitopes, did not induce T cell-mediated cytotoxicity (Fig 5a). However, cells infected with Ad5-MCMV_IE1, which express the immunodominant MCMV IE1 protein, were selectively killed by T cells from MCMV-infected mice and not from MCMV-naïve counterparts (Fig 5b). Some residual cytotoxicity was detected in the MCMV-naïve group and may be explained by the expression of stress ligands by the mSGM cells, which activates T cell-mediated cytotoxicity, or by secretion of cytokines from T cells incidentally activated by the purification process [22]. To control for differential cell growth rates, apoptosis is shown as a ratio of individual fluorescent cells to percent confluence of the monolayer. Apoptosis in Ad5-MCMV_IE1-infected mSGM cells with MCMV-experienced T cells increased significantly from the 24-hour time point to the 48-hour time point (P<0.01; comparison not shown on figure). Fluorescent events were gated by size to remove background fluorescence and any T cells which underwent apoptosis (detailed in *Materials and Methods* section).

**Figure 5.**
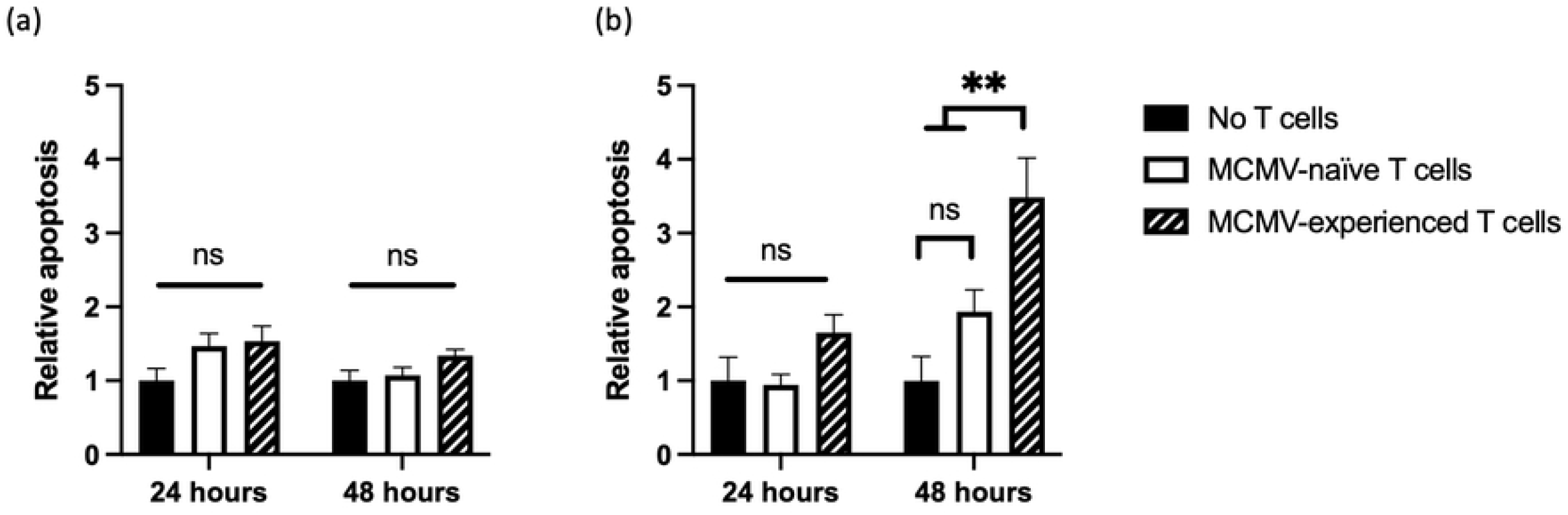
mSGM cells are vulnerable to *ex vivo* MCMV antigen-specific T cell mediated cytotoxicity. mSGM cells were infected with *Ad5-control* (a) or *Ad5-MCMV_IE1* (b). Apoptosis is shown as a ratio of apoptotic cells compared to time 0. Cells alone (black) or MACS-enriched splenic T cells from uninfected (white bar) or MCMV-infected (lined bar) mice at 28 days post infection were added at E:T ratio of 6:1. Data represent 2 animals with 6 technical replicates each (total n=12).

### mSGM cells are suitable for lentivirus transduction

Retroviral vector transduction is a valuable tool for constructing stable transgene-expressing cell lines, and we therefore investigated whether mSGM cells could be readily transduced using a lentivirus. To test this the MCMV m164 protein, which is one of two known immunodominant T cell epitopes in BALB/c mice during MCMV latency, was transduced into mSGM cells [23]. A control lentivirus expressing enhanced green fluorescent protein (eGFP) and mCherry and a second lentivirus expressing a FLAG-tagged MCMV m164 protein under an HCMV promoter were evaluated. Transduction efficiency reached nearly 100% in the control cells, with all visible cells positive for mCherry. After selection with puromycin, m164-expressing cells (mSGM-m164) were compared with un-transduced cells and did not appear to differ morphologically (Fig 6a-b). Cellular m164 expression could be confirmed by staining with anti-FLAG antibodies (Fig 6c). These results further demonstrate the broad potential applicability of the mSGM cell line.

**Figure 6.**
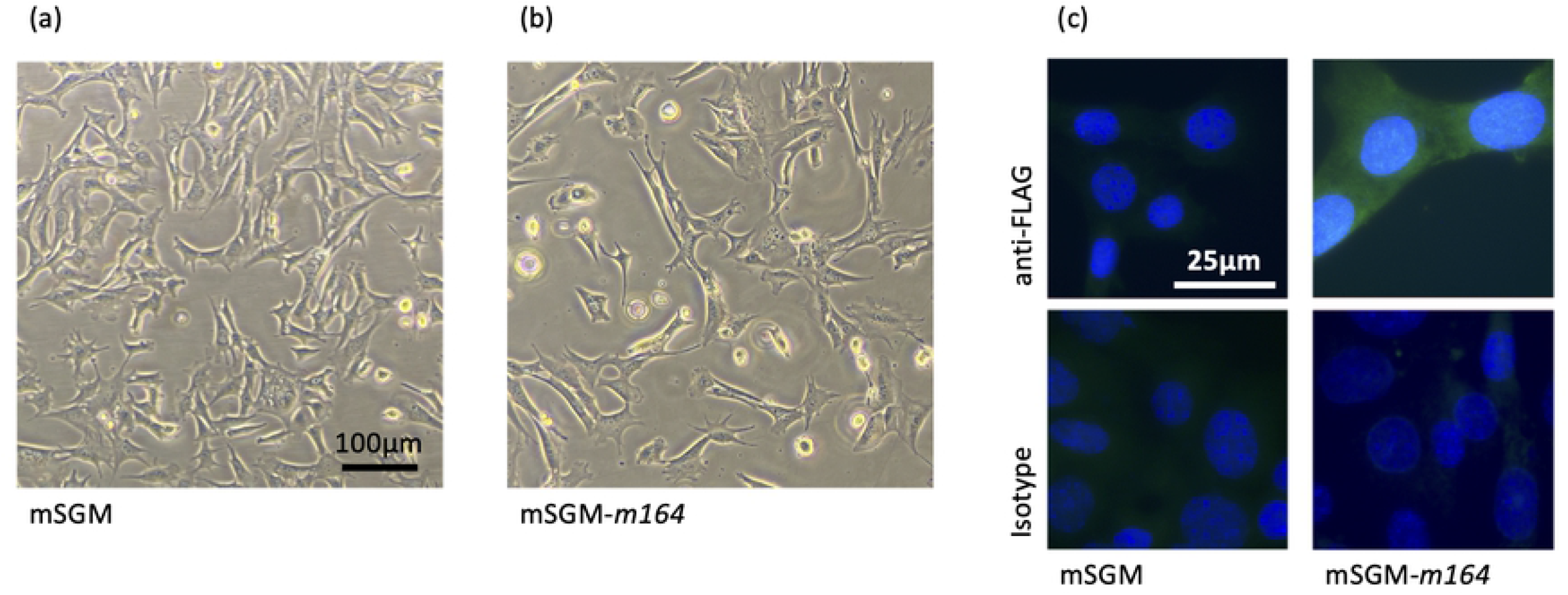
mSGM cells are suitable for lentivirus transduction. (a-b) mSGM cells were imaged before (a) and after transduction (b) with MCMV m164-FLAG lentivirus. (c) After selection, cells were stained for FLAG epitope (upper panels) or isotype control (lower panels) and counterstained with DAPI.

## Discussion

MCMV infection of BALB/c mice is considered an important animal model for human cytomegalovirus infection. The mouse immune response to MCMV infection is similar in many ways to the human immune response to HCMV, and the tissue and cell tropism and replication kinetics of both viruses follow a similar pattern [24]. BALB/c mice are also inexpensive and easily maintained in comparison with larger CMV host species, such as guinea pigs or rhesus macaques. However, despite the benefits of the BALB/c mouse model, commercially-available cell lines used to produce MCMV virus stocks and for plaque assays are typically NIH 3T3 cells, a mouse embryonic fibroblast cell line isolated from a Swiss albino mouse. NIH 3T3 cells therefore express different cellular markers, including different MHC haplotypes. This precludes their use in some common *ex vivo* assays such as T cell- or NK cell-mediated cytotoxicity assays. To remedy this issue, many researchers opt to derive their own mouse embryonic fibroblasts (MEFs), which can be a laborious process. The stable mSGM cell line described herein may provide a useful alternative.

High expression of all 3 MHC class I isotypes on mSGM cells allows presentation of MCMV peptides, providing a powerful, easily maintained system for *ex vivo* cytotoxicity assays. The m164- and IE1-derived immunodominant peptides are presented on H-2D^d^ and H-2L^d^ MHC isotypes, respectively [23]. Our studies did not confirm precisely which epitopes were presented, only that antigen-specific T cell-mediated cytotoxicity could be detected in response to both m164 and IE1 proteins. Our flow cytometry results indicate that all 3 isotypes are present on the cell surface. Future investigation of MHC peptide complexes on transgene-expressing or virus-infected mSGM cells could be used to confirm epitopes matching those seen in MCMV-infected BALB/c mice. Transgene-expressing mSGM cells were readily produced using simple lentivirus transduction, extending their possible uses. In particular, the m164 protein which was efficiently expressed after lentivirus transduction is an important T cell target in MCMV-infected BALB/c mice. A similarly transduced line may therefore be useful for CTL assays in the future.

In addition to the BALB/c strain-specific proteins that mSGM cells express, the mSGM cell line was also found to be significantly more sensitive during plaque assays. Detection of low numbers of viral particles in sera, tissue homogenates, or culture supernatant is subject to the limit of detection of the assay. Quantitative PCR is a common substitute for plaque assays, with very high sensitivity, but the presence of viral genome copies does not always correlate with infectious viral particles. Using mSGM cells, it was found that infectious MCMV could be detected with 3- to 4-fold greater sensitivity compared to the same sample titrated on NIH 3T3 fibroblasts. Centrifugal enhancement, a method commonly employed to detect low concentrations of viral particles, and previously demonstrated for MCMV [11], increased the sensitivity 100- to 150-fold over standard plaque assay on NIH 3T3 cells. This increase in sensitivity may be particularly useful when detecting virus in samples where infectious virions are found in very low numbers.

Taken together, these findings suggest that the mSGM cell line provides substantial improvements in sensitivity of plaque assays and *ex vivo* T cell cytotoxicity assays for studies of MCMV. The ease of culture, hardiness in a variety of culture medium, and tolerance of freeze-thaw cycles, as well as the stable morphology at low and high passage, all emphasize the usefulness of this cell line. mSGM cells may therefore shed insight on cytomegalovirus pathogenesis and immune responses and be useful for other infectious disease models.

## Materials and Methods

### Culture and selection media

NIH 3T3 cells (American Type Culture Collection, Manassas, VA, USA; ATCC #CRL-1658) and primary salivary gland cells were maintained in complete medium consisting of Dulbecco’s modified Eagle medium with 4.5 g/l glucose and L-glutamine without sodium pyruvate (DMEM; Corning, NY, USA) supplemented with 10% v/v fetal bovine serum (FBS; Hyclone, Logan, UT, USA), 0.3% sodium bicarbonate, 4 mM HEPES (Gibco, Waltham, MA, USA), 2 mM L-glutamine (Gibco), and 1x gentamycin solution (Gibco). Selection of SV40 large T antigen-positive cells was performed using complete medium described above, with the addition of 250 µg/ml G418 (Geneticin; Gibco). Selection of m164 and control lentivirus-transformed cells included the addition of 2 µg/ml puromycin (Gibco). After immortalization, salivary gland cells were found to grow more rapidly and FBS was reduced to 4-8% v/v to prevent overgrowth. 293A cells (ThermoFisher, Waltham, MA, USA; catalog #R70507) and 293FT cells (ThermoFisher; catalog #R70007) were grown in DMEM supplemented with 10% FBS and 1x penicillin/streptomycin (Gibco).

### Isolation of salivary gland cells

Submandibular salivary gland tissue was collected from specific pathogen-free 6-week-old female BALB/c ByJ mice obtained from Jackson Laboratory (Bar Harbor, ME, USA). Salivary gland tissues were minced with scissors into roughly 2 mm^3^ pieces and cultured in complete medium in a 6-well tissue culture plate (Corning, NY, USA) until cells began to form a monolayer. When the monolayer approached confluency, the tissue fragments were removed and the monolayer was dissociated with 0.25% trypsin containing EDTA (Gibco, Waltham, MA, USA). The resulting cell suspension was filtered through a 40 µm cell strainer to remove any remaining tissue fragments. Cells were then re-plated at a density of 3 × 10^5^ cells per well in 6-well plates for transfection. After plating, cells were allowed 24 hours to become completely adherent.

### Transfection with SV40 large T antigen

The pSV3-neo plasmid expressing the SV40 large T antigen and a neomycin resistance cassette (American Type Culture Collection, Manassas, VA, USA; ATCC# 37150) was prepared and purified using ZymoPURE Plasmid Miniprep Kit (Zymo Research, Irvine, CA, USA) according to manufacturer’s instructions. 1µg of purified pSV3-neo was diluted in OptiMEM medium (Gibco), to which diluted Lipofectamine 2000 (Invitrogen, Waltham, MA, USA) in OptiMEM was added at a 1:1 ratio. The solution was incubated on ice for 1 hour. For transfections, 6-well culture plates containing sub-confluent monolayers of primary salivary gland cells were used. Culture medium was removed and 300µl plasmid transfection solution was added per well. Cells were incubated for 3 hours at 37° C. Then, transfection solution was removed and replaced with 3 ml fresh culture medium. After 24 hours, roughly 50% cell death had occurred and medium was changed to remove debris. At 48 hours post-transfection, G418 selection agent was added to the medium at a concentration of 100µg/ml. An un-transfected control well was used to ensure that the G418 concentration was enough to induce cell death. After the initial selection phase, the G418 concentration was reduced to 50µg/ml for maintenance. Medium was changed every 3-5 days.

### Viruses and virus stock preparation

K181-Perth strain MCMV was kindly provided by Helen Farrell at the University of Queensland. Virus stocks were prepared as previously described in NIH 3T3 cells [7,10]. Virus stock titer was determined by plaque assay on NIH 3T3 cells as described below.

Plaque assays were conducted as previously described [7]. Briefly, samples were diluted serially at 10-fold increments in complete medium, and 500µl of each dilution was incubated on a 60-70% confluent monolayer of NIH 3T3 or mSGM cells in a 12-well culture plate at 37° C. After 1 hour, medium was removed and wells were overlaid using 0.75% w/v carboxymethylcellulose (CMC; ThermoFisher, Waltham, MA, USA) in modified Eagle medium (MEM; Gibco) supplemented to a final concentration of 10% FBS, 2mM L-glutamine, 0.5x non-essential amino acid solution (NEAA; Gibco), and 1x gentamycin. At 4-5 days post infection, wells were washed, fixed with 100% methanol, and stained with modified Giemsa stain (Sigma-Aldrich, St. Louis, MO, USA). Selection agents were not added to plaque assay overlay medium due to the terminal nature of the assay. For centrifugal enhancement, plates were centrifuged at 250*g* for 1 hour at 4°C in place of the 1-hour incubation [11].

### Replication assays

Low- and high-MOI growth curves were performed using mSGM cells or NIH 3T3 cells as previously described [12,13]. Briefly, 12-well tissue culture plates were seeded with mSGM or NIH 3T3 cells and incubated in complete medium with the indicated MOI of MCMV for 1 hour. Medium was then removed and replaced with fresh complete medium and cultures were incubated for 6 days. At 1-day intervals, wells were scraped using a cell scraper and cells and supernatant were collected and frozen. Collected samples were thawed and sonicated, and titer was determined by plaque assay on NIH 3T3 cells as described above.

### Animal infection studies

4- to 6-week-old female BALB/c ByJ mice were obtained from Jackson Laboratory (Bar Harbor, Maine, USA) and maintained under specific pathogen free conditions at the Louisiana State University Department of Laboratory Animal Medicine. For infection, mice were injected intraperitoneally with 10^6^ PFU MCMV in 0.2 ml PBS. Mock-infected animals were injected with 0.2 ml PBS. At 60 days post infection, mice were euthanized with 2.5% Avertin followed by exsanguination. Tissues were collected and kept on ice until use. All animal protocols were approved by the Louisiana State University Institutional Animal Care and Use Committee.

### Flow cytometry

Monolayers of mSGM cells were dissociated with 0.25% trypsin-EDTA (Gibco). Cells were then washed twice with phosphate buffered saline containing 2% FBS, passed through a 70-µm cell strainer, and fixed with 2% formaldehyde. Extracellular staining was conducted with the following antibodies: αH-2L^d^-PE, αH-2D^d^-AF647, αH-2K^d^-AF488 (BioLegend, San Diego, CA), and matched isotype controls. Flow cytometry was performed immediately after staining using a Foressa X-20 flow cytometer (Beckton Dickinson). Results were analyzed with FlowJo software (Beckton Dickinson).

### Immunofluorescence staining

For immunofluorescence imaging, cells were grown to 70% confluence on 8-well glass culture slides (Merck Millipore, Tullagreen, Carrigtwohill, Ireland), washed, fixed with 2% formaldehyde, and washed again in PBS. Primary antibodies were added at 1:100 dilutions and slides were incubated at 4° on a rocker overnight. Cells were then washed thoroughly with PBS and secondary antibodies were added at a 1:100 dilution for 1 hour. After washing with PBS, slides were cover-slipped using ProLong Diamond Antifade Mountant with DAPI (Invitrogen, Eugene, OR) and glass coverslips. Primary antibodies were rat αCD31, αCD140a, and αCD324 (Beckton Dickinson) and isotype control. The secondary antibody used was chicken anti-rat IgG AF488 (Invitrogen).

### T cell enrichment

Whole mouse spleens were homogenized in 3 ml complete medium and filtered through 70-micron cell strainers. Nucleated cells were separated from red blood cells and cell debris using Lympholyte M (Cedarlane Laboratories Ltd., Canada) according to manufacturer’s instructions. Cells were then pelleted by centrifugation and counted using a hemocytometer. T cells were enriched from spleen homogenate by magnetic-activated cell sorting (MACS) using Miltenyi Pan T Cell Isolation Kit (Miltenyi Biotec, Bergisch Gladbach, Germany) according to manufacturer’s instructions, using an input of 10^7^ cells. Enrichment was confirmed to be >95% efficient by flow cytometry. T cells were resuspended in RPMI 1640 (Gibco) with 10% FBS, 1x penicillin/streptomycin solution (Gibco), and beta-mercaptoethanol.

### Live cell apoptosis assay

mSGM or mSGM-*m164* cells were added to a 96-well flat-bottomed cell culture plate at a density of 1×10^4^ cells per well. After allowing 12-24 hours for cells to adhere, culture medium was replaced with RPMI containing 10% FBS and 1x penicillin/streptomycin solution (Gibco). MACS-enriched T cells were then added at the indicated effector:target ratio. NucView 488 fluorescent caspase-3/7 substrate (Biotium, Fremont, CA, USA) was added at a final concentration of 1.25µg/ml to visualize cells undergoing apoptosis. Live-cell imaging was conducted using an Incucyte system set to scan 2 fields from each well of a 96-well cell culture plate at 1-hour intervals. In an initial study, effector:target ratios were tested from 1:1 to 10:1. The ratio 6:1 was found to provide optimal T cell-mediated killing with minimal background fluorescence during the 48-hour test period. After 48 hours, experiments were ended due to an increase in background fluorescence and monolayer overgrowth.

### Cloning

Cloning was performed using the FastCloning method described by Li et al. [14]. The MCMV IE1 gene was cloned into the plasmid pAd5-Blue [15] to make pAd5-MCMV_IE1. First, PCR amplification of the IE1 gene was conducted using 10ng of purified MCMV (KP) DNA. Primers ClaI_MCMV_IE1_F 5’-AGAGAGATCGATCACCATGGAGCCCGCCGCAC-3’ and SpeI_FLAG-MCMV_IE1_R 5’-AGAGAGACTAGTCTACTTATCGTCGTCATCCTTGTAAATCCTT-3’ were used to amplify the IE1 gene. The PCR product was then gel purified on a 1% agarose gel and visualized with ethidium bromide. DNA was extracted using the Zymoclean gel DNA recovery kit (Zymo Research, Irvine, CA). 1 μg of purified IE1 DNA was then digested with ClaI and SpeI restriction enzymes (New England Biolabs, Ipswich, MA). Digested DNA was cleaned using the DNA clean and concentrator kit (Zymo Research, Irvine, CA). 1 μg pAd5-Blue was digested with ClaI and XbaI and dephosphorylated using Quick CIP (New England Biolabs, Ipswich, MA). Enzymes were heat inactivated by heating to 80 °C for 5 minutes and DNA precipitated using 100% ethanol. Digested pAd5-Blue and IE1 PCR products were then ligated using T4 DNA ligase (New England Biolabs, Ipswich, MA) according to manufacturer’s instructions. Plasmids were then transformed into XL-10 Gold chemically competent *Escherichia coli* cells (Agilent Technologies, Santa Carla, CA) as previously described [14]. The resulting plasmid was designated pAd5-MCMV_IE1.

For lentivirus plasmid generation, first the MCMV m164 gene was PCR amplified from 10 ng purified MCMV DNA. Primers were pLV_m164_FC_F 5’-AAGCAGGCTGCCACCATGTTTCTCCGCGGCTGTCG-3’ pLV_m164-FLAG_R 5’-ACAAGAAAGCTGGGTCTACTTATCGTCGCATCCTTGTAATCAGATGAACGTTCGACG CGC-3’. The m164 PCR product was then purified as above and ligated into the plasmid pLV (Vector Builder, Chicago, IL). For control lentivirus, insert was amplified using primers pLV_CDS_FC_F 5’-ACCCAGCTTTCTTGTACAAAGTGGTG-3’ and pLV_CDS_StartFC_R 5’-GGTGGCAGCCTGCTTTTTTGTAC-3’. The ligated products were then transformed into XL-10 Gold competent cells as previously described [14].

### MCMV-antigen-expressing adenovirus and lentivirus vector production

Adenovirus containing the MCMV IE1 gene was generated by transfecting purified pAd5-MCMV_IE1 into 293A cells in 1 well of a 6 well plate using Lipofectamine 3000 (ThermoFisher, Waltham, MA) according to manufacturer’s instruction. Once plaques were visualized, the plate was freeze/thawed 3 times and cellular debris pelleted by centrifugation at 1000g for 5 minutes. 100μL of supernatant was then used to infect a 70% confluent T-75 flask of 293A cells and infection allowed to progress for 5 days. Following infection, cells were freeze thawed 3 times and cellular debris pelleted by centrifugation at 1000 x g for 5 minutes. The resulting supernatant was then titrated on 293A cells by limiting dilution and frozen at -80 °C until use.

Lentivirus containing the MCMV m164 gene was produced by co-transfection of pLV-m164-FLAG with psPAX2 (a gift from Didier Trono, Addgene plasmid #12260) and pCMV-VSV-G (a gift from Bob Weinberg, Addgene plasmid #8454) [16] mixed at a 1:1:1 molar ratio using Lipofectamine 3000 into 293FT cells (ThermoFisher, Waltham, MA). Supernatant was collected after 4 days and frozen at -80° C until use. For transduction of mSGM cells, lentivirus was added directly to culture medium of sub-confluent monolayers of mSGM cells in 12-well tissue culture plates. After 48 hours, cells were selected under 1 ng/mL puromycin. Control wells without lentivirus were used to ensure that the concentration of puromycin was sufficient to induce 100% cell death in the absence of resistance.

### Data analysis and visualization

Data were analyzed using Prism 9 (GraphPad, San Diego, USA). For most figures, data were tested for normality and ANOVA with Tukey’s post-hoc test was used. Graphs were produced using Prism 9. Flow cytometry data were analyzed and visualized with FlowJo (Beckton Dickinson). Live cell images were analyzed with Incucyte software with parameters set to exclude fluorescent cells smaller than 300 µm^2^, which allowed us to exclude any apoptotic leukocytes (which are typically 9-11 µm in diameter, or under 100 µm^2^) while including mSGM cells (which are closer to 30 µm in diameter, or 700 µm^2^ during apoptosis).

## Author contributions

TW designed and carried out experiments and wrote manuscript. BS designed and assisted with experiments, produced lentivirus and adenovirus, and edited manuscript. CB assisted with experiments and edited manuscript. JR provided SV40 plasmid and carried out transfection. RC designed experiments, provided funding, and edited manuscript.

## Acknowledgments

This work is supported by NSTP fellowship grant number AP19VSNL00C037 from the USDA Animal and Plant Health Inspection Service. Assistance with flow cytometry was provided by Marilyn Dietrich in the Louisiana State University flow cytometry core facility.

